# FLASHQuant: a fast algorithm for proteoform quantification in top-down proteomics

**DOI:** 10.1101/2023.11.08.566181

**Authors:** Jihyung Kim, Kyowon Jeong, Philipp T. Kaulich, Konrad Winkels, Andreas Tholey, Oliver Kohlbacher

## Abstract

Liquid chromatography-mass spectrometry (LC-MS) based top-down proteomics (TDP) is an essential method for the analysis of intact proteoforms. The accurate quantification of individual proteoforms is a crucial step in identifying proteome-wide alterations in different biological conditions. Label-free quantification (LFQ) is the most common method for proteoform quantification as it requires no additional costly labeling. In TDP, due to frequent co-elution and complex signal structures, overlapping signals deriving from multiple proteoforms complicate accurate quantification. Here, we introduce FLASHQuant for MS1-level LFQ analysis in TDP, which is capable of automatically resolving and quantifying co-eluting proteoforms. FLASHQuant performs highly accurate and reproducible quantification in short runtimes of just a few minutes per LC-MS run. To validate the proteoforms reported by FLASHQuant, we evaluated them with identified proteoforms confirmed by tandem mass spectrometry, which showed high match rates. FLASHQuant is publicly available as platform-independent open-source software at https://openms.org/flashquant/.

## INTRODUCTION

Top-down proteomics (TDP) based on mass spectrometry (MS) is an emerging analytical approach for the comprehensive identification and characterization of intact proteoforms^1–6^. The term proteoform refers to all of the different molecular forms of a protein that can originate from variations at genetic or protein levels, i.e. posttranslational modifications^7^. TDP’s advantage over the conventional bottom-up approach is its ability to preserve the intact form of proteins by skipping the enzymatic digestion step prior to MS, thus providing a more complete and accurate representation of the biologically relevant proteoforms. In the past decade, research in TDP has seen substantial improvements in protein separation, MS techniques, and bioinformatic software and demonstrated its potential to elucidate the important role of proteoforms in biomedical processes^3,8–10^. Technical advances have enabled researchers to move forward from qualitative analysis to quantitative analysis on proteoform studies^11–13^.

Quantitative analysis is crucial in proteomics, as it opens the door for comparative studies of proteins in different biological functions or for biomarker discovery^11,14,15^. To relatively quantify proteoform abundances, in principle, three general strategies can be employed: label-free quantitation (LFQ), metabolic labeling, and chemical labeling. Metabolic labeling has seldom been employed in TDP, mainly due to inherent challenges such as interfering isotope patterns between light/heavy labeled proteins^16–18^. In contrast, chemical labeling-based strategies such as isobaric labeling have been successfully applied^19,20^ but still need further improvements regarding the labeling procedures as well as the bioinformatics data interpretation.

Therefore, the most widely applied approach in TDP is LFQ^12,13,21^, which performs a relative quantification by direct comparison between MS runs, which provides advantages such as low costs (no need for expensive labeling reagents), fewer experimental steps, and omission of increased sample complexity (i.e., introduced in chemical labeling approaches due to incomplete or over-labeling).

Nevertheless, the analysis of TDP-LFQ data still is a major bottleneck. Existing LFQ data analysis tools for MS1 level quantification (including chromatographic feature detection and intensity calculation) still need further developments and improvements regarding reproducibility, overlapping signal resolution, or usability.

LFQ of proteoforms in most cases still relies on deconvolution-centric software, such as Protein Deconvolution (Thermo Fisher Scientific)^22,23^, DataAnalysis (Bruker Daltonics)^24–26^, TopPIC suite^6,27,28^, and ProMex^29,30^. As the primary goal of deconvolution tools is to detect proteoforms’ masses rather than quantify them, chromatographic information needed for quantification is widely neglected on these platforms. This results in limited peak coverage and compromises reproducibility in quantification. Therefore, many LFQ studies have been performed using in-house scripts to retrieve chromatographic data based on deconvoluted proteoform masses.

Despite its significance, the problem of overlapping signals in LFQ studies has often been overlooked^2,31^. Overlapping signals occur when the distinct proteoforms interfere with each other (i.e., sharing mass-to-charge ratio (*m/z*) values), primarily due to the complexity of the samples and the co-elution of multiple proteoforms from the LC within a narrow retention window. TDP, which deals with larger analytes compared to bottom-up proteomics or metabolomics, exacerbates this issue; the larger size of analytes (proteins/proteoforms) leads to broader isotopic envelopes and wider charge state ranges, compared to those of peptides, contributing to the dataset’s high complexity. As a result, co-eluted proteoforms of different charge states can occupy similar or overlapping *m/z* ranges even when they have highly distinct masses^2,32^, introducing possible quantification bias in particular in the above-mentioned deconvolution-centric approaches. Thus, the overlapping signal issue should be addressed for accurate LFQ in TDP, which necessitates advanced data analysis strategies.

Not only have minimal TDP-LFQ data analysis software been developed, but most of them have not been freely released to users. QMT^21^ and IPQuant^33^ have been introduced and utilized but are not publicly available. Commercial platforms such as ProSightPD (Proteinaceous, Inc., Evanston, USA) have been widely used but, unfortunately, are not freely available.

Here, we introduce FLASHQuant, a fast and robust tool for MS1-level LFQ analysis in TDP. FLASHQuant incorporates an individual chromatogram extraction procedure and a conflicting feature (i.e., overlapping proteoform signals) resolution method using a non-negative least squares solver. In our evaluation, FLASHQuant demonstrated its effectiveness compared to the widely used ProSightPD. As part of OpenMS, FLASHQuant is publicly available as platform-independent open-source software with a graphical user interface at https://OpenMS.org/FLASHQuant.

## RESULTS

### Structure of FLASHQuant

FLASHQuant is composed of two main stages: feature group detection and quantity calculation (Figure 1). Feature group refers to the group of LC features of different charge states from a single putative proteoform and its isotopes. For accurate quantification, the detected feature groups are refined while the quantities are calculated.

**Figure 1.**
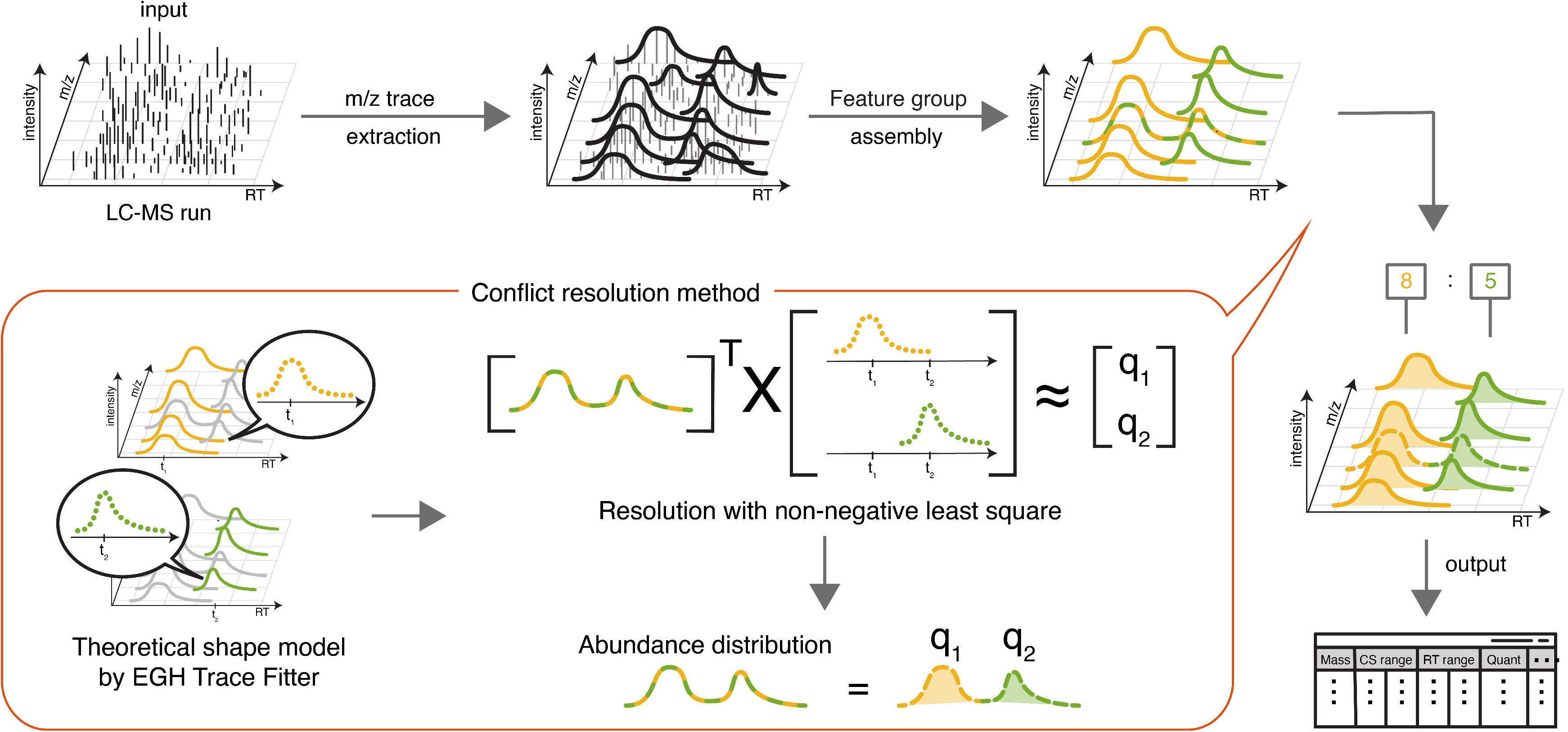
Illustration of FLASHQuant algorithm. From a centroided LC-MS1 full scan, FLASHQuant extracts *m/z* traces (individual ion chromatograms), denoted as black curved lines at the top-center. Detected *m/z* traces are then assembled into feature groups, representing putative proteoforms. Feature groups (depicted as yellow and green) often share *m/z* traces (the mixed-colored line), which should be resolved for accurate quantification. The theoretical shapes of each feature group are estimated using its non-shared *m/z* traces by fitting them against the EGH function. Based on the estimated shapes (having retention time and intensity values), the proportion of each shape (q1 and q2) in the shared trace is estimated with a non-negative least square solver. Then, the shared *m/z* trace quantity is distributed between the corresponding feature groups. The feature group abundances are finally calculated as the summed area of *m/z* traces in each feature group. The output of FLASHQuant is given as a tsv format file by default.

The feature group detection stage starts with extracting *m/z* traces (individual ion chromatograms) from the LC-MS spectra using MassTraceExtractor^34^. MassTraceExtractor is a self-consistent kernel density estimator to connect peaks having *m/z* values within the tolerance (defined by the instrument’s mass accuracy) along the retention time. Thus, each extracted *m/z* trace represents a chromatographic signal having a specific charge state and isotopic composition over an elution window. The extracted *m/z* traces are deconvolved and assembled into feature groups by FLASHDeconv^35^. Both charge and isotope deconvolution are applied to *m/z* traces to determine the masses. In this step, an *m/z* trace may be assigned to multiple feature groups representing the overlapping signal (the shared *m/z* trace) from distinct proteoform ions (the feature groups).

In the next stage, the abundance of feature groups is calculated while resolving conflicting feature groups. FLASHQuant starts by resolving shared *m/z* traces for accurate quantity calculation (similar to EPIQ^36^) in the conflict resolution method. The shared *m/z* traces are mainly from overlapping signals but also from errors in feature group detection, where an error in feature group detection arises when an *m/z* trace is assigned to an incorrect feature group. While different in origins, both cases can be addressed by comparing the *m/z* trace elution profiles within the feature groups containing the shared *m/z* trace (termed conflicting feature groups). For each shared *m/z* trace, FLASHQuant constructs theoretical shapes (the representative *m/z* trace shapes) of its conflicting feature groups. The theoretical shape of a feature group is generated by selecting its non-overlapping *m/z* traces and fitting them against the exponential-Gaussian hybrid function^37^. On generating the theoretical shapes for the conflicting feature groups, FLASHQuant attempts to assemble the shared *m/z* trace with the theoretical shapes by solving a non-negative least square problem. While solving, the proportion of each theoretical shape in the shared trace is estimated, which can be used to allot the abundance of the shared *m/z* trace to each of the conflicting feature groups. After all shared *m/z* traces are resolved, the abundance of each feature group is calculated by summing areas under all corresponding *m/z* trace curves (see STAR Methods for details of each step).

Using centroided MS1 data as input, FLASHQuant outputs information on feature groups, including their monoisotopic masses, retention time range, charge range, and quantities (see Table S1 or S2). Additionally, we provide an extra tool, ConsensusFeatureGroupDetection, allowing users to easily attain jointly detected feature groups among multiple LC-MS runs (i.e., technical replicates) within mass and retention time tolerance.

### Dataset description

We evaluated the performance of FLASHQuant using three datasets with different complexity: **PIPMix**, **SpikeIn**, and **ProteomeMix**. PIPMix is the least complex dataset consisting of single LC-MS runs with a six-protein mixture (Pierce Intact Protein Standard Mix). To generate the SpikeIn dataset, different amounts of the PIPMix (relatively diluted 1, 1/2, 1/3, 1/5, 1/7, and 1/10 with 20 ng/µl as 1) were spiked into an *E. coli* lysate, each of which was subject to an LC-MS run. The ProteomeMix dataset contains varying proportions of proteins from a human Caco-2 cell lysate (1/5, 1/2, 1, 2, and 5) with a constant concentration for *E. coli* lysate.

Prior to MS analysis, proteoforms were separated with a 90-minute linear gradient on an Ultimate 3000 nano-UHPLC system equipped with a reversed-phase C4 column coupled online to a Fusion Lumos Tribrid mass spectrometer. MS1 spectra were acquired at high resolution (120,000), and within a cycle time of 4 s, the most intense ions were fragmented by collision-induced dissociation.

We benchmarked FLASHQuant against ProSightPD (“ProSightPD Hi Res. Feature Detector” node, version 4.2 in Proteome Discoverer 3.0, Thermo Fisher Scientific) in all comparisons (see STAR Methods for the details on parameters). In terms of runtimes, FLASHQuant achieved a strong advantage over ProSightPD; for the PIPMix and SpikeIn datasets together, FLASHQuant took a total of 44 minutes, whereas ProSightPD took a total of 17 hours and 42 minutes. For the ProteomeMix dataset, FLASHQuant took 19 minutes, and ProSightPD took 4 hours and 14 minutes.

### FLASHQuant and ProSightPD achieved high sensitivity with the SpikeIn dataset

As a first validity check on proteoform detection and quantification sensitivity, we compared the performance between FLASHQuant and ProSightPD with the SpikeIn dataset. Figure 2A shows the quantification result of four spiked-in proteins, as separately depicted in four columns, from all dilutions and technical replicates in the SpikeIn dataset (see Table S1 for the full lists of results for both FLASHQuant and ProSightPD). Due to the chosen MS setting, only four spiked-in proteins of mass below 28 kDa were detected. Both tools succeeded in detecting all four proteins across dilution samples (down to a 1/10 ratio) and triplicates, proving their limit of detection ranges. However, FLASHQuant showed higher quantification linearity, as fold changes from ProSightPD are more dispersed from the expected values than FLASHQuant. This dispersion is more evident for the larger proteins (21 and 28 kDa) with lower abundance compared to smaller proteins (9 and 11 kDa). For the 21 kDa protein, average differences in fold change from the expected value were 0.06 and 0.14 for FLASHQuant and ProSightPD, respectively, and 0.06 and 0.16 for the 28 kDa protein.

**Figure 2.**
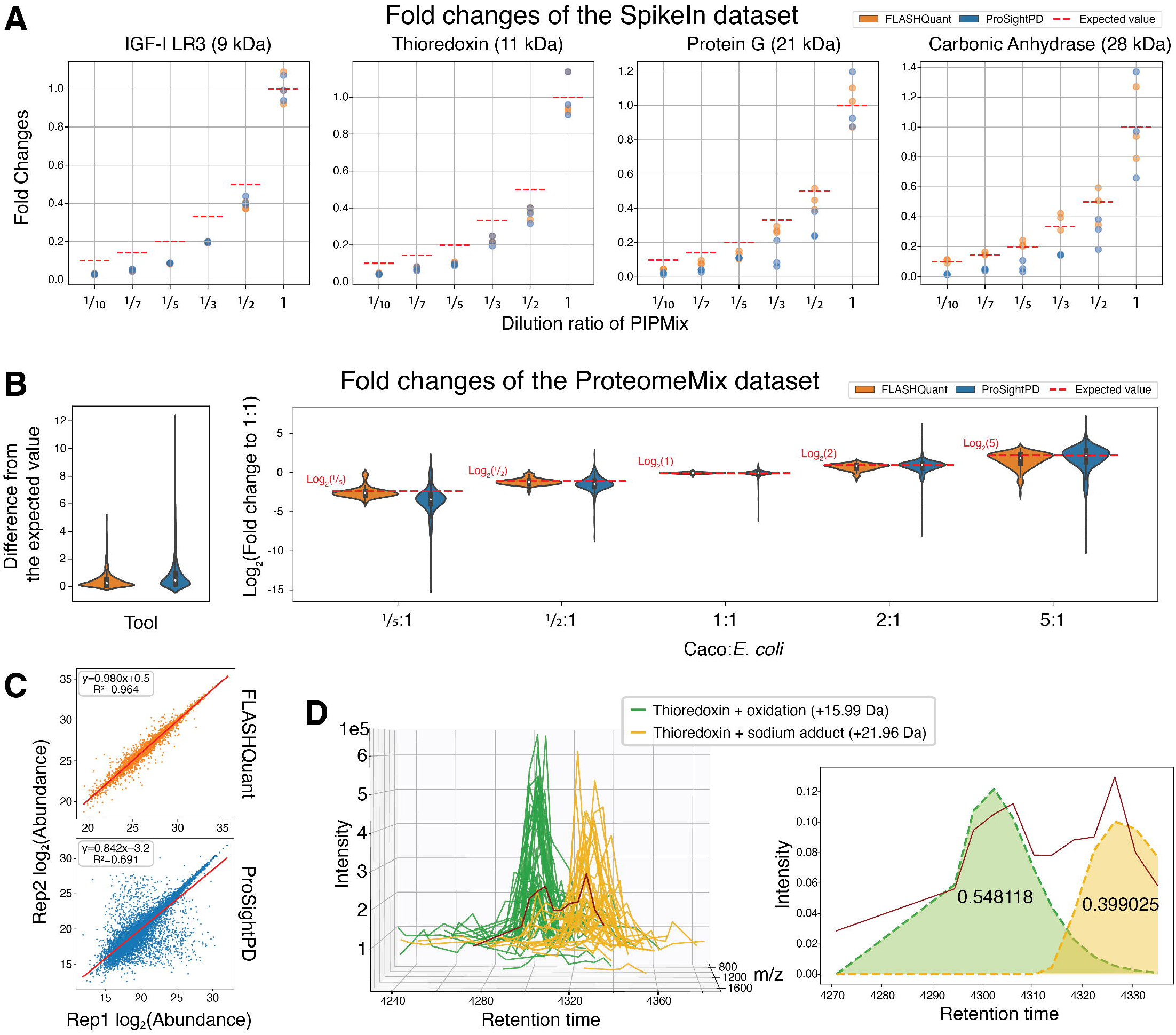
Data analysis of the SpikeIn, ProteomeMix, and PIPMix datasets. A. The fold changes comparison for the SpikeIn dataset. The four spiked-in proteins are shown in four columns, with the expected fold change values as red dashed lines. The x-axis in each column displays three pairs of dots (per two colors), representing fold change values from three replicates. These fold changes were computed based on the average quantity of spiked-in protein at 20 ng/µl concentration, denoted as “1.” FLASHQuant showed unbiased quantification across all samples, demonstrating its high quantification accuracy and reproducibility. B. The fold changes comparison for the ProteomeMix dataset. The consensus feature groups of all samples (5 different mixing ratios and two replicates each) that were matched to the human Caco-2 proteoforms (identified by TopPIC) were employed in this analysis. The violin plot on the left shows the fold change difference from the expected values, and the figure on the right depicts the logarithmized fold change values per mix ratio. FLASHQuant shows less variance to the expected values and its closer mode values to the expected values. See Figure S5 for the analysis substituted TopPIC with ProSightPD Search. C. Quantification reproducibility comparison between replicates for the ProteomeMix dataset. The linear regression between each quantity of the jointly detected feature groups was depicted as a red line. FLASHQuant demonstrated its high reproducibility with a higher R² value and regression slope closer to 1 than ProSightPD. See Figure S4 and S6 for only the detected feature groups matched against human Caco-2 proteoforms. D. An example of how the conflict resolution method works in FLASHQuant. Two proteoforms from the SpikeIn dataset with a 6 Da mass difference were indicated as green and yellow. One of their shared *m/z* traces is colored in dark red as an example. The left figure shows all raw *m/z* traces of both proteoforms, and the figure on the right illustrates the modeled theoretical trace shapes for each proteoform. The black digits on the theoretical shapes refer to the resolved abundance ratio for each proteoform.

Moreover, when we calculated the coefficient of variation (CV) of replicates with feature group quantity, FLASHQuant reported low median CV values of ∼0.15 from the four proteins, showing high reproducibility, while ProSightPD delivered higher values (∼0.28) (Figure S1). Similar to the fold change analysis, FLASHQuant showed its strength in reproducibility for the larger proteins by delivering smaller CV values.

### FLASHQuant delivers a strong connection to the identification

In order to demonstrate a detailed understanding of the reported feature groups, we attempted to explain the masses of the output feature groups from the simplest dataset available, PIPMix. This is achieved by identifying proteoforms from the MS2 spectra in the dataset and comparing their masses to the feature groups reported by FLASHQuant and ProSightPD (see Table S1 for the full list of feature groups).

We began by detecting, for each tool, consensus feature groups among all output feature groups from technical triplicates. The default parameter settings of ProSightPD, a mass tolerance of 100 ppm and a retention time tolerance of 8 min (used for all consensus feature group detection from hereon) were applied to this process. For the detection of consensus feature groups in FLASHQuant, we utilized the ConsensusFeatureGroupDetection. The masses of consensus feature groups were then compared to the proteoform masses identified by either TopPIC or ProSightPD Search within 20 ppm mass tolerance (see Table S3 for the full list of identified proteoforms and Table S5 for the detailed number of identified proteoforms). ProSightPD Search was separate from the ProSightPD LFQ (ProSightPD Feature Detector) and consisted of “ProSightPD 4.2 Annotated Proteoform Search” node and “ProSightPD 4.2 Subsequence Search” node. It was separately executed to analyze MS2 spectra (see STAR Methods for details on parameters).

Figure S2 shows the Sankey diagrams of each tool’s consensus feature groups, comparing those matched against identified masses within 20 ppm mass tolerance to those that did not match. Half of the FLASHQuant results matched identified proteoform masses compared to 38.2% for ProSightPD, demonstrating the high connectivity of FLASHQuant with the identification results.

Among these consensus feature groups, unidentified consensus feature groups that did not match identified proteoform masses do not necessarily imply false positives. Instead, some of them showed a potential for modified proteoforms that were not detected by database searches.

We took unidentified consensus feature groups and compared them to the masses of the identified proteoforms allowing one PTM mass (±500 Da and has Unimod accession number). For FLASHQuant, 32.2% of the consensus feature groups were explained by this comparison, while 43.6% for ProSightPD (See Table S4 for details). 16.4% for both FLASHQuant and ProSightPD resulted in unexplained. Since the detection of feature groups on the MS1 level does not necessarily translate to identifications on the MS2 level (i.e., insufficient fragmentation or database search limitations), a large portion of unexplained feature groups is not surprising. Those from FLASHQuant had plausible raw signals (see Figure S3 for examples), indicating a possibility of multiple PTMs or truncation.

### Performance on proteome-wide quantification

FLASHQuant performs rather conservative feature group detection and quantification, leading to fewer consensus feature groups than ProSightPD at the proteome level (348 and 768 in the ProteomeMix dataset, respectively. See Table S2 for the full list of feature groups). However, the quantification variances from FLASHQuant were relatively small, while its quantification reproducibility was high (See Figure 2B and Figure 2C). Also, when we matched the masses of those consensus feature groups against the identified proteoform masses from TopPIC, 82.2% were matched for FLASHQuant while 67.6% for ProSightPD. These are the same trends as the above analysis of the PIPMix dataset, with more consensus feature groups from FLASHQuant being validated by identified proteoforms than from ProSightPD.

To test the quantification accuracy, we evaluated fold changes of human Caco-2 proteoforms in the ProteomeMix dataset (Figure 2B). Among the reported feature groups from each LC-MS run, those with a mass within 20 ppm tolerance to identified human Caco-2 proteoform masses were selected for further analysis. The average difference between the measured and expected fold changes is smaller in FLASHQuant (Figure 2B left panel), with median values of 0.24 and 0.49. The right panel of Figure 2B shows that the values of all ratio samples in FLASHQuant are closer to the expected values with smaller variance than in ProSightPD, proving its high quantification linearity as in SpikeIn dataset analysis.

Quantification reproducibility between replicates was demonstrated through a linear regression between each quantity of the jointly detected feature groups (Figure 2C). Between the replicates of each sample, we collected the jointly detected feature groups (same mass and retention time tolerance as consensus feature groups) and then evaluated the similarity of their quantities. FLASHQuant showed high reproducibility with R² values of 0.98 and regression slopes of 0.96, while ProSightPD resulted in less concentrated values to the linear regression line, with R² values of 0.84 and regression slopes of 0.69.

When only the consensus feature groups matching the identified proteoforms by TopPIC were considered (Figure S4), both tools showed high reproducibility with R² values of >0.9 and regression slopes of 1±0.1. However, slightly closer-to-one R² values were observed in FLASHQuant than ProSightPD in all samples.

Additionally, the same fold changes and reproducibility analyses, except for substituting TopPIC with ProSightPD Search, also confirmed a similar outcome (see Figures S5 and S6).

### Resolving overlapping proteoforms boosts the quantification accuracy

FLASHQuant experienced a boost in both SpikeIn and ProteomeMix datasets using the conflict resolution method. When FLASHQuant was executed without the conflict resolution method (“resolving-off-mode” from hereon vs. “default-mode”), still a large portion of reported feature groups (82-85% for SpikeIn and 74-80 % for ProteomeMix) overlapped with the default-mode. These overlapped feature groups (within mass and retention time tolerances) from the two modes even showed similar dynamic ranges. The exclusive ones from the resolving-off-mode were mostly eliminated during or after the conflict resolution method in the default-mode, as most of their *m/z* traces were incorrectly assigned to them (Figure S7).

The conflict resolution method not only removed possible errors in the feature group assignment but also alleviated the quantification errors. From the ProteomeMix dataset, the exclusive ones from the resolving-off-mode have larger fold change differences from the expected values compared to the default-mode (Figure S8).

A detailed example of how the conflict resolution method performed on proteoforms is illustrated in Figure 2D, which shows the raw signals (i.e., *m/z* traces) from two proteoforms of the target protein Thioredoxin (11,858.04 Da) in the SpikeIn dataset (ratio ⅓). The green traces are Thioredoxin with an oxidation (+15.99 Da), while the yellow with a sodium adduct (+21.96 Da). Due to a difference of only 6 Da in mass between the two proteoforms, they inevitably shared some *m/z* traces, depicted as dark red. However, with the conflict resolution method, the abundance of the shared signal was successfully distributed to the two proteoforms, with 54.8% attributed to the oxidized proteoform and 39.9% to the proteoform with the sodium adduct (the remaining 5.3% corresponds to the portion unexplained by both proteoform feature groups, e.g., noisy component).

### FLASHQuantWizard: GUI for running FLASHQuant

We provide a graphic user interface (GUI), FLASHQuantWizard, for users to conveniently run FLASHQuant (Figure S9). The development of FLASHQuantWizard was derived from SwathWizard in OpenMS. SwathWizard is an effective pipeline GUI tool that assists SWATH proteomics data analysis. Some of the SwathWizard’s useful segments, such as LC-MS file loader and displaying logs, are shared with FLASHQuantWizard.

The main segment of FLASHQuantWizard contains three widgets; Output selection, Output consensus feature group, and Run FLASHQuant. In the first widget, the Output selection widget, users can specify an output directory or ask for additional output files in featureXML format. Also, the next widget, the Output consensus feature group widget, offers additional consensus feature group output files (from here on, consensus files) among replicates upon request. When the input files contain replicates generated under multiple experimental conditions, each requiring separate consensus files, FLASHQuantWizard can distinguish these replicates. For this, the shared substring between replicates’ file names should be given by users. Otherwise, all input files are used to output a consensus file. The last widget, Run FLASHQuant, allows for parameter adjustment of FLASHQuant.

## DISCUSSION

We here introduce FLASHQuant, a robust quantification tool designed explicitly for MS1-level LFQ data analysis in TDP. One of the key features of FLASHQuant is its automatic conflicting resolution method, which addresses the commonly overlooked yet critical problem of overlapping signals. By effectively resolving overlapping signals, FLASHQuant ensures highly accurate quantification and offers remarkable reproducibility among technical replicates. Benefiting from the ultrafast and robust FLASHDeconv algorithm, FLASHQuant achieves a rapid runtime of ∼2 min per dataset (containing over 1800 MS1 spectra).

FLASHQuant demonstrated its advantage in high accuracy and reproducibility in quantification. It is showcased by unbiased quantification across a wide range of samples, exemplified by its impartial quantification of all four proteins within the SpikeIn dataset. In contrast, ProSightPD yielded biased results as more significant fold change and the CV value differences were observed in larger proteins (21 kDa and 28 kDa) with lower abundances, while smaller proteins exhibited fewer variations. This observed disparity in outcomes may be attributed to the distinctive quantification methodologies employed by the two tools. FLASHQuant initiates the analysis by detecting elution profiles (i.e., *m/z* traces) and subsequently deconvolving them, while ProSightPD follows a different approach by deconvolving the masses before connecting them with the corresponding traces.

In addition to FLASHQuant, we also provided two accompanying tools, ConsensusFeatureGroupDetection and FLASHQuantWizard, to enhance the user experience. We have implemented a Python script to visualize the overlapping signals, such as in Figure 2D. In the near future, we plan to develop a simple web application that provides interactive visualization of quantified feature groups (which will be shared at https://openms.org/flashquant/). This application will greatly assist users in visually examining the raw signals, which is essential in LFQ. Moreover, given the strong reliance of FLASHQuant on FLASHDeconv and both tools being part of OpenMS, we are actively working on integrating the results from both tools, which will offer users a comprehensive approach that combines the strengths of both tools.

It is worth noting that while FLASHQuant excels in accuracy and reproducibility, ProSightPD outperforms it in terms of sensitivity. This difference can be attributed to FLASHQuant’s use of strict scoring thresholds, resulting in a lower number of feature groups compared to ProSightPD. These stringent thresholds were implemented to achieve high specificity in the analysis and, in turn, delivered high-quality results (i.e., high connectivity of the feature groups to the proteoform identifications).

Furthermore, in order to further enhance the reproducibility of FLASHQuant, the addition of a retention time alignment method could be applied. This alignment method would reduce variation in retention times across technical replicates, thereby improving the consistency and comparability of quantification results. Additionally, we acknowledge the importance of incorporating statistical functions in conjunction with FLASHQuant results. To address this need, we are exploring the possibility of integrating statistical methods directly into FLASHQuant or providing users with R scripts for performing comprehensive downstream statistical analysis on the FLASHQuant results.

## ACKNOWLEDGEMENTS

J.K., K.J., and O.K. were funded by Horizon 2020 Marie Sklodowska-Curie Action ITN 2017 of the European Commission (grant 765502-A4B). K.J. and O.K. acknowledge EPIC-XS (project number 823839), funded by the Horizon 2020 program of the European Union. A.T. is funded by the DFG Cluster of Excellence “Precision Medicine in Inflammation”

## AUTHOR CONTRIBUTIONS

O.K. conceived the idea of the proteoform LFQ algorithm for the TDP MS datasets and K.J. proposed the conflict resolution method. J.K. and K.J. developed and implemented FLASHQuant algorithm. P.T.K., K.W., and A.T. designed the experiment and performed the sample preparation and LC-MS experiments. All authors analyzed the data. J.K., K.J., and P.T.K. wrote the manuscript with input from all authors, which was read, commented on, and approved by all authors.

## STAR METHODS

### RESOURCES AVAILABILITY

#### Lead contact

Further information and requests for resources and reagents should be directed to and will be fulfilled by the lead contact, Jihyung Kim.

#### Materials Availability

This study did not generate new unique reagents.

#### Data and Code Availability

- The mass spectrometry proteomics data have been deposited at the ProteomeXchange Consortium via the MassIVE repository and are publicly available. The accession number and DOI are listed in the key resources table.
- FLASHQuant original code has been deposited and is publicly available at https://github.com/JeeH-K/OpenMS/tree/feature/FLASHQuant
- Any additional information required to reanalyze the data reported in this paper is available from the lead contact upon request.

## EXPERIMENTAL MODEL AND SUBJECT DETAILS

### Human Caucasian colon adenocarcinoma cultivation

The cultivation of human Caucasian colon adenocarcinoma (Caco-2) cells was maintained as per European Collection of Authenticated Cell Cultures recommendation. The cells were grown at 37 °C with 5% CO_2_ in RPMI-1640 medium (25 mM HEPES, 2 mM L-glutamine, 13 nM phenol red) supplemented with 10% (v/v) fetal bovine serum and 1% (v/v) penicillin (10,000 U/ml). After reaching 90-100% confluence, the cells were passaged using TrypLE™ Express enzymes to detach the cells. Prior to harvesting, the cells were washed three times with PBS buffer (centrifugation at 200×g for 5 min at 25 °C). The cells were stored at −80 °C prior to cell lysis.

### Escherichia coli cultivation

*Escherichia coli* (strain MG1655) was cultured at 37 °C in M9 minimal medium containing 15 mM glucose, as described previously^38^. In brief, bacterial cells were grown to reach an OD_600_ of 1. The culture was divided into 50 ml aliquots and centrifuged (3,000×*g*, 5 min, 25 °C) to pellet the cells. The cells were washed twice in MilliQ water and stored at −80 °C prior to cell lysis.

## METHOD DETAILS

### Methanol-Chloroform-Water precipitation

Methanol-Chloroform-Water precipitation was performed according to Wessel & Flügge^39^. In brief, 150 µl of the sample was mixed with 600 µl methanol, 150 µl chloroform, and 450 µl MilliQ water. The mixture was centrifuged (14,000×*g*, 20 min, 25 °C) and the upper phase was removed. 600 µl of methanol was added, mixed thoroughly, and centrifuged. The supernatant was removed, and the protein pellet was washed twice with 600 µl methanol prior to air drying.

### Generation of the ProteomeMix sample

E. *coli* cells and Caco-2 cells were lysed in 1% sodium dodecyl sulfate, 10 mM TRIS (pH 8.8), 1× cOmplete protease inhibitor (Promega, Madison, USA) by ultrasonication and protein concentration was determined by the Pierce BCA protein assay kit (Thermo Fisher Scientific, Bremen, Germany). Approximately 500 µg of proteins were purified by methanol-chloroform-water precipitation and subjected to GELFrEE fractionation (8% tris-acetate cartridge) according to manufacturer’s protocol (Expedeon, Eching, Germany). In brief, the samples were mixed with 30 µl 5× sample buffer, 8 µl 1 M dithiothreitol, 112 µl MilliQ and incubated 10 min at 50 °C (1,400 rpm) prior to fractionation. To enrich proteoforms smaller than approximately 30 kDa, the first GELFrEE fraction was used from the Caco-2 or *E. coli* sample, respectively^40^. The fractions were purified by chloroform-methanol-water precipitation, and the proteins were dissolved in MS loading buffer (3% acetonitrile, 0.1% trifluoroacetic acid (TFA)). The Caco-2 and *E. coli* samples were analyzed by LC-MS/MS to determine total ion counts (TICs) and were diluted with MS loading buffer to yield approximately the same intensity of TICs.

In order to obtain the proteome mixture, Caco-2 and *E. coli* proteins were mixed in five different ratios (from 1:5 to 5:1), keeping the concentration of *E. coli* proteins constant and varying only the amount of Caco-2 proteins (Table S6).

### Generation of the SpikeIn sample

For the generation of the SpikeIn sample, small *E. coli* proteins (<20 kDa) were enriched by solid-phase extraction, as described previously^41^. In brief, *E. coli* cells were lysed in 8 M guanidium hydrochloride, 1× cOmplete protease inhibitor (Promega) by freeze-thaw cycling. The sample was heated 10 min at 70 °C and acidified with 5% formic acid (FA). After centrifugation (20 min, 21,100×*g*, 4 °C), the supernatant was transferred to an activated (methanol) and equilibrated (5% FA) C18 solid-phase extraction cartridge (3 cc 200 mg Waters, Eschborn, Germany). The proteins were washed twice with 5% FA prior to elution of the small proteins with 70% and 100% acetonitrile. The proteins were vacuum-dried and resuspended in MS loading buffer. Protein concentration was determined by the Pierce BCA protein assay kit (Thermo Fisher Scientific).

To obtain the SpikeIn samples, the Pierce™ Intact Protein Standard Mix (Thermo Fisher Scientific, six proteins ranging from 9-68 kDa (human IGF-I LR3 (P05019, 40-118): 9,105.3482 Da; human Thioredoxin (Q99757, 60-166): 11,858.04393 Da; *Streptococcus dysgalactiae* Protein G (P06654, 223-413): 21,429.75915 Da; bovine carbonic anhydrase (P00921, full length): 28,963.6881 Da; *Streptococcus* Protein AG (P02976, P19909, chimeric): 50,429.84641 Da; *Escherichia coli* Exo Klenow (P00582, 324-928) 67,959.42515 Da), resuspended in MS loading buffer) was added to the small protein enriched *E. coli* lysate. The concentration of *E. coli* proteins was kept constant (160 ng/µl), while the final concentration of the protein mixture was varied (20 ng/µl, 10 ng/µl, 6.67 ng/µl, 4 ng/µl, 2.86 ng/µl or 2 ng/µl).

### LC-MS/MS analysis

Proteoforms were separated using an Ultimate 3000 nano-UHPLC system (Thermo Fisher Scientific) equipped with an analytical reversed-phase C4 column (Accucore, 50 cm×75 μm, 2.6 μm, 150 Å, Thermo Fisher Scientific). A precolumn (C4 PepMap 300, 5 µm, 300 Å, Thermo Fisher Scientific) in forward-flush mode was used for sample loading. Eluent A was 0.05% FA and eluent B was 80% acetonitrile, 0.04% FA. The separation was performed over a 90 min gradient from 15-55% B: 0-5 min 4% B, 5-7 min 4-15% B, 7-97 min 15-55% B, 97-99 min 55-98% B, 99-110 min 98% B, 110-110.1 min 98-4% B, 110.1-120 min 4% B.

The UHPLC system was coupled online to a Fusion Lumos Tribrid mass spectrometer (Thermo Fisher Scientific). MS1 spectra (400-1800 *m/z*) were acquired with a resolution of 120,000, 246 ms maximum injection time, 200% normalized AGC target, and 4 microscans. Within a cycle time of 4 s, the most intense ions were selected (charge states: 4-50 and undetermined charge states with dynamic exclusion enabled (n=2, 60 s) for fragmentation with collision-induced dissociation (25%). The settings for MS2 spectra were: 60,000 resolution, 250 ms maximum injection time, 1,000% normalized AGC target. The acquisition was performed in intact protein mode (ion-routing multipole pressure of 2 mTorr) and lock-mass was enabled (445.12003 *m/z*). The RF value was set to 30% and source-induced dissociation was 15 V.

### *m/z* trace extraction

The *m/z* trace extraction is performed using two algorithms in OpenMS, MassTraceDetection and ElutionPeakDetection^34^. MassTraceDetection takes centroid LC-MS spectra as input and connects the peaks within *m/z* tolerance that continuously appear across the retention time. FLASHQuant takes five ppm for this *m/z* tolerance as a default value to only gather peaks from the same analyte (of distinct charge state and isotope) into a single *m/z* trace. Since distances between isotopes from an intact protein with a large charge state are relatively smaller than those from peptide or metabolite, a small *m/z* deviation should be employed to distinguish those isotopes. After MassTraceDetection effectively separates and groups spectral peaks from distinct compounds with different *m/z* values, ElutionPeakDetection further separates them along the retention time direction by analyzing the shape of each *m/z* trace. For each *m/z* trace, its abundance is determined by its total area, and its retention time range is truncated with its full width at half maximum. This truncation is done since the elution profile often holds long tails with low intensities. Note that different *m/z* traces may represent the same (proteoform) molecule but with different isotopes or charges here. Grouping the *m/z* traces from a putative single molecule is done in the next feature group assembly step.

### Feature group assembly

In this step, detected *m/z* traces from each putative proteoform are assembled into a feature group using a fast and robust spectral deconvolution algorithm FLASHDeconv. During its deconvolution to find monoisotopic masses, FLASHDeconv automatically groups the spectral peaks from different masses. Since the core algorithm of FLASHDeconv takes a spectrum as input, we first convert the *m/z* traces into a series of spectra along the retention time direction. To convert *m/z* traces along retention time into spectra, we first bin the retention time with a fixed bin size of the minimum retention time span of the *m/z* traces. Then, for each bin, all *m/z* traces whose retention time range overlaps with the bin are first collected. Then, the collected traces are projected into a single spectrum, where peak intensities are determined by the abundances of the projecting traces. If different bins share exactly the same set of *m/z* traces, only one bin out of them was processed to avoid redundancy.

Each generated spectrum is then deconvolved using the FLASHDeconv algorithm so that *m/z* traces are collected into feature groups. Each feature group has its (monoisotopic) mass, abundance (the summation of its member *m/z* traces), and cosine score (a fit score to the theoretical isotope patterns) determined by FLASHDeconv. The retention time range of a feature group is given by the union of the ranges of its member *m/z* traces. While exactly the same spectra are not generated from the above conversion, still many (in particular, the ones from consecutive bins) would yield the same or quite similar feature groups that are likely to be from the same proteoform. Next, we attempt to detect such feature groups and merge them into a single one. To this end, first, the feature groups are sorted in descending order of their abundances. Then, we iterate over the sorted feature groups (denoted by *f*) and perform the following merging process i) - iv):

i) For each iteration for the feature group *f*, all the feature groups whose masses are within 10 Da from the mass of *f* and whose abundances are less than the abundance of *f* are selected.
ii) Then only the ones whose retention time ranges overlap with that of *f* by more than 50% are retained. The remaining feature groups are denoted by *F^−^*.
iii) From *F^−^*, we further discard the feature groups *f*’ such that the mass difference between *f*’ and *f* is larger than 3 Da and no more than 50% of the *m/z* traces in *f*’ overlap with those in *f*. Note that *F^−^* contains *f* itself.
iv) Generate a merged feature group *f^−^* by taking all *m/z* traces in feature groups within *F^−^*. FLASHDeconv is again employed to calculate the mass, abundance, and cosine score for *f^−^*. Only if the cosine score of *f^−^* is higher than *f^−^*, *f* is replaced by *f* in the original feature group set *F^−^*. Otherwise, *f* is unchanged (and thus, the above steps have no effect).

After this merging process is done for all features in *F*, the conflict resolution is performed as described below.

### Conflict resolution method

The conflict resolution method aims to separate shared *m/z* traces of different feature groups in the feature group set *F*. The generated feature groups often share *m/z* traces with each other due to co-elution or simply mass artifacts. This conflict should be resolved by distributing the quantity of *m/z* trace to the shared feature groups. Therefore, this stage only concerns feature groups in conflict with others. Moreover, since sharing of the *m/z* traces between different feature groups is highly localized, we divided the *m/z* traces in each feature group into subgroups (called features) with the same charge states and used the features as the basic unit for the conflict resolution. Since each feature consists of the *m/z* traces from the same (putative) mass of the same charge, they are highly localized in the *m/z* dimension.

For fast conflict resolution, it is important to quickly detect conflicting features (i.e., features sharing the same *m/z* traces) and find clusters of shared features. Veit^42^ suggested the idea of these conflicts being modeled as an undirected graph *G* = (*V*, *E*), where nodes v in V represent features and two nodes v and v’ are connected by an edge when their representing features have a conflict. By finding the edge-connected subgraphs on this graph, one can quickly find the clusters of conflicting features.

For each cluster, the conflict resolution method is applied for each edge of the cluster, similar to the scheme already established and proved in BUP^36^. The key idea of this scheme is to reconstruct shared *m/z* traces using the unshared *m/z* traces. In theory, all the *m/z* traces’ elution profiles in a feature (and further those in a feature group) should be the same, and the traces shared by different features should have the shape of a weighted summation of the unshared traces from the conflicting features. To get the theoretical elution profile of a feature (from hereon, theoretical shape), first, the unshared traces are collected from each feature. Then, the theoretical shape is modeled based on an exponential-Gaussian hybrid function^37^ using the EGHTraceFitter algorithm in OpenMS. If no unshared *m/z* trace exists for a feature, we track the feature group that contains the feature, find the most abundant *m/z* trace in that feature group, and use it to model the theoretical shape.

Upon finding the theoretical shapes from the conflicting features, the conflict resolution can be formulated by a distribution problem in which we want to distribute the quantity of the shared *m/z* trace to the features. This distribution problem can be described as a non-negative least square problem.

Let *M* be a matrix composed of theoretical shapes *A*,

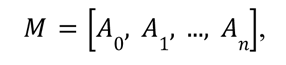

where *n* is the number of theoretical shapes, thus the number of corresponding features. *Y* denotes the shape of the shared *m/z* trace, and the lengths of each *A*s are adjusted to match the length of *Y* by zero padding or truncation.

We compute a vector *Q* of length *n*, *q* ∈ *Q*, that solves

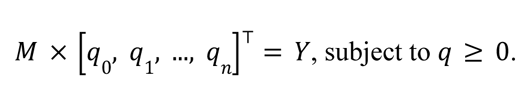

*q* refers to the proportions of features to describe the quantity of shared *m/z* trace. Each *q* is multiplied by the quantity of the *m/z* trace and distributed over the corresponding features.

### Tool parameters

We used FLASHQuant with its default parameters, including the *m/z* tolerance of 10 ppm, the mass tolerance of 3 Da, and the isotope cosine threshold of 0.85. Only mass and charge ranges were modified for each dataset based on the characteristic of the target analyte; the mass range and the charge range were set to 1-70 kDa and 2-100, respectively, for the SpikeIn and the PIPMix datasets, and 1-30 kDa and 2-50 for the ProteomeMix dataset. Since the Pierce Intact Protein Standard Mix from Thermo Fisher includes a protein with a mass of 68 kDa, we decided to broaden the mass and charge ranges for SpikeIn and PIPMix datasets. Likewise, ProSightPD was used with the default “LFQ processing method” parameter settings, except for the mass and charge ranges. The mass and charge ranges were consistent with FLASHQuant.

TopPIC (version 1.6.4) and ProSightPD Search were utilized to analyze MS2 spectra from all datasets and to identify proteoforms’ masses. Again, both tools are executed with their default parameters except for the mass and charge ranges, consistent with FLASHQuant.

### Filtering feature groups to the masses of identified proteoforms

With the PIPMix dataset, we aimed to explain all the reported feature groups as much as possible. Thus, we first conducted an open search with TopPIC to gather a possible modification list. In total, 12 modifications were chosen based on the most frequent mass shifts (Disulfide, Dehydro, Didehydro, Amidated, Pro->pyro-Glu, Trp->Oxolactone, Methyl+Deamidated, and Dioxidation) and 4 common modifications suggested by TopPIC (Acetyl, Phopho, Oxidation, and Methyl). Using these 12 modifications as variable modifications, TopPIC was executed once more to collect identified proteoform masses. The six protein sequences given by Thermo Fisher were taken as a database. For ProSightPD Search, variable modifications needed to be annotated per site on each sequence. Therefore, instead of using the FASTA format database, the XML format database equipped with a modification list was downloaded from UniProt. As some of the UniProt proteins had a slightly different sequence compared to the protein sequences given by Thermo Fisher, we utilized the “Database Manager” to modify the input sequences manually.

The database for the ProteomeMix dataset was generated by combining the *E. coli* database (Swiss-Prot, taxon ID 83333, downloaded from UniProt in July 2023) with the human database (Swiss-Prot, taxon ID 9606, downloaded from UniProt in July 2023). The dataset and database for ProteomeMix are far larger than PIPMix, so TopPIC could not be run twice. Also, database search with variable modifications was extremely time-consuming (more than a day for one file), such that only an open search was performed. XML format of the equivalent database (also including modifications) was downloaded from UniProt and was employed for ProSightPD Search.

## SUPPLEMENTAL INFORMATION

**Figure S1.** Coefficient of variation (CV) values from technical replicates of the SpikeIn dataset

**Figure S2.** Sankey diagram of the number of consensus feature groups from the PIPMix dataset

**Figure S3.** Examples of raw mass traces of FLASHQuant consensus feature group from the PIPMix dataset that did not match against identified masses

**Figure S4.** Reproducibility of FLASHQuant and ProSightPD consensus feature groups matched against TopPIC identified masses from the ProteomeMix dataset

**Figure S5.** Analogue of Figure 2B for the consensus feature group masses matched against Human CaCo-2 proteoform masses detected by ProSightPD Identification

**Figure S6.** Analogue of Figure S4 for the consensus feature groups matched against ProSightPD identification

**Figure S7.** Barplot of resolving-off-mode exclusives analysis

**Figure S8.** Boxplot of fold change differences from the expected values comparing FLASHQuant exclusives to resolving-off-mode exclusives with the ProteomeMix dataset

**Figure S9.** FLASHQuantWizard

**Table S1.** The outputs from FLASHQuant and ProSightPD for the SpikeIn and PIPMix datasets

**Table S2.** The outputs from FLASHQuant and ProSightPD for the ProteomeMix dataset

**Table S3.** The identification outputs from TopPIC and ProSightPD Search for the PIPMix and ProteomeMix datasets

**Table S4.** The consensus feature group outputs from FLASHQuant and ProSightPD for the PIPMix dataset and their matches to the identification results

**Table S5.** The number of identified proteoforms by ProSightPD Search and TopPIC for the PIPMix dataset

**Table S6.** Generation of the ProteomeMix

