## Supplemental materials for "FLASHQuant: a fast algorithm for proteoform quantification in top-down proteomics"

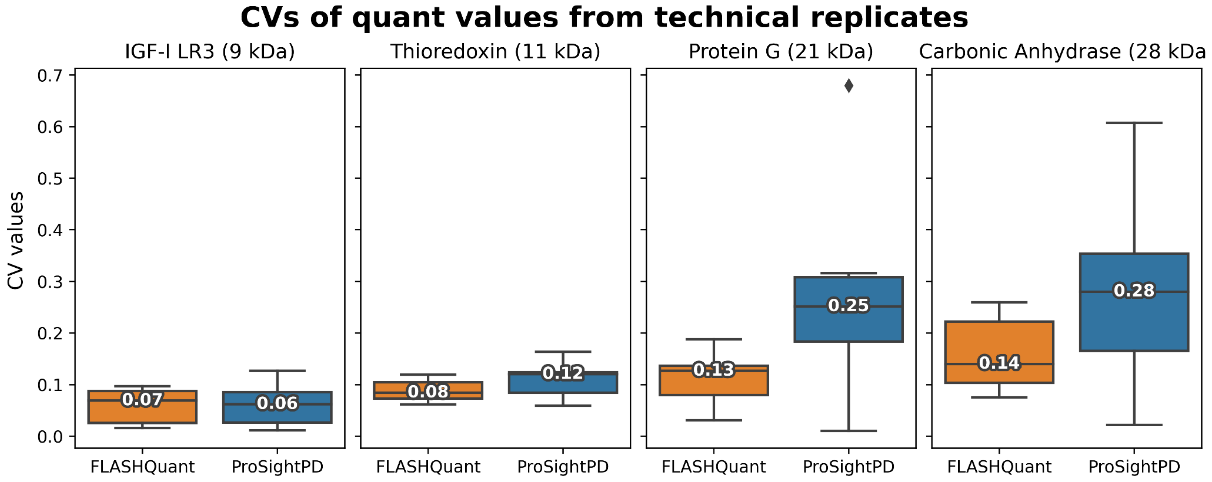


**Figure S1. Coefficient of variation (CV) values from technical replicates of the SpikeIn dataset**

Boxplot of CV values calculated from quantification values of technical replicates per spiked-in proteins. The digits on the boxplot indicate their median CV values. FLASHQuant shows consistently low CV values from all four proteins, while ProSightPD results in high CV values in larger molecules. Note that Carbonic Anhydrase is not only the largest protein but also has the lowest abundance among the four proteins.

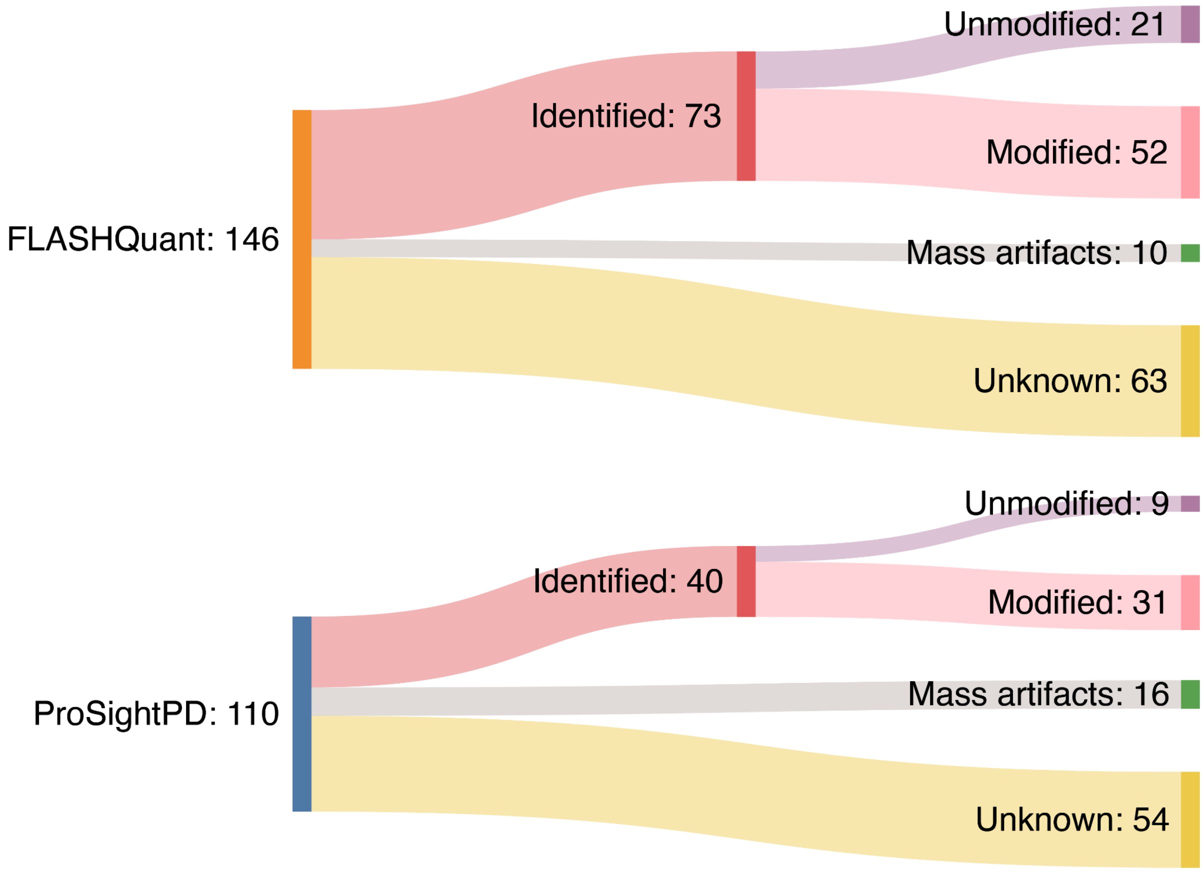


**Figure S2. Sankey diagram of the number of consensus feature groups from the PIPMix dataset**

A Sankey diagram shows the number and type of consensus feature groups. Identified type refers to consensus feature groups having the same mass as identified proteoforms and has two flows depending on whether they matched unmodified or modified identified proteoforms. Mass artifacts type includes isotopologues and harmonics. Half of the FLASHQuant results belonged to the Identified type, whereas half of the ProSightPD results were of the unknown type, confirming FLASHQuant's strong connection to the identification results.

(a) Proteoform with the mass 3526.99 Da

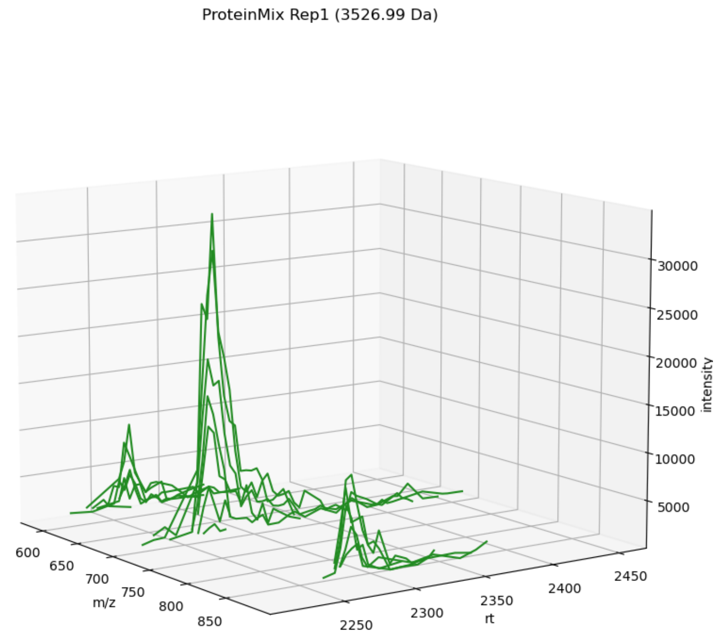


(b) Proteoform with the mass 15497.49 Da

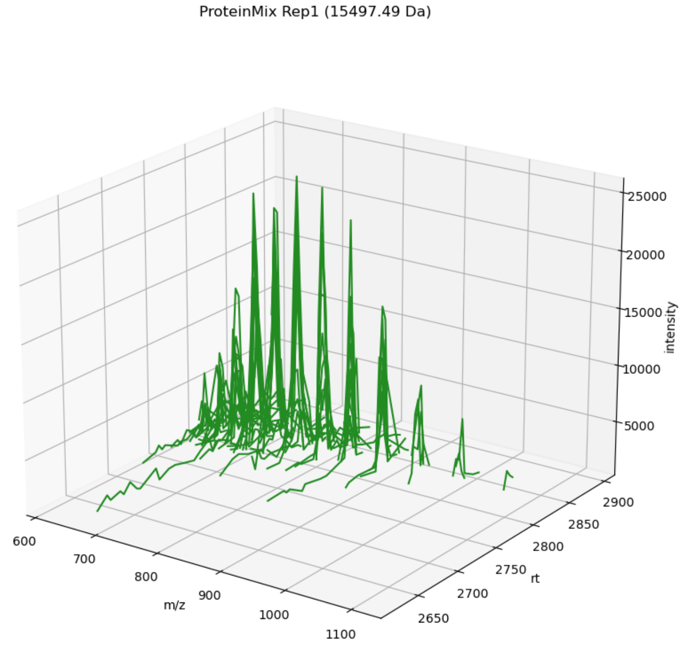


**Figure S3. Examples of raw mass traces of FLASHQuant consensus feature group from the PIPMix dataset that did not match against identified masses**

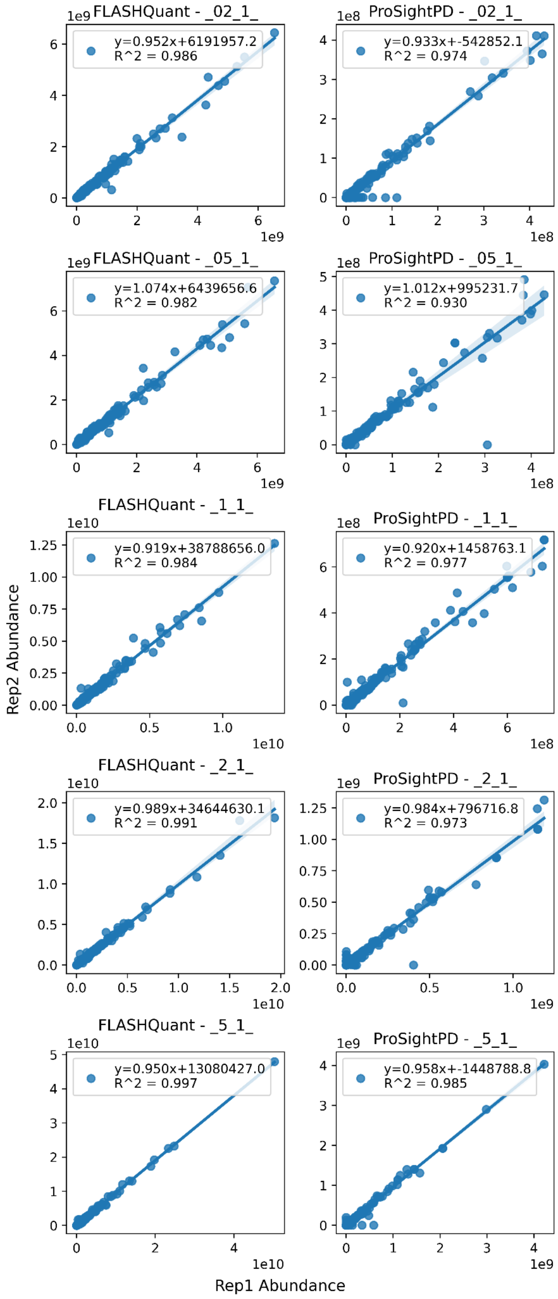


**Figure S4. Reproducibility of FLASHQuant and ProSightPD consensus feature groups matched against TopPIC identified masses from the ProteomeMix dataset**

Reproducibility between technical replicates has been evaluated by calculating linear regression between each quantity of the consensus feature groups. Results from FLASHQuant are in the left column, and the ones from ProSightPD are in the right column. Each row indicates five different samples from the ProteomeMix dataset (1/5,1/2, 1, 2, and 5, from top to bottom). Feature group quantities from the first replicate are on the x-axis, for the second replicate on the y-axis. Regression slopes and R2 values are written in the left corner, showing high reproducibility from both tools with regression slopes of 1±0.1 and R2 values of >0.9.

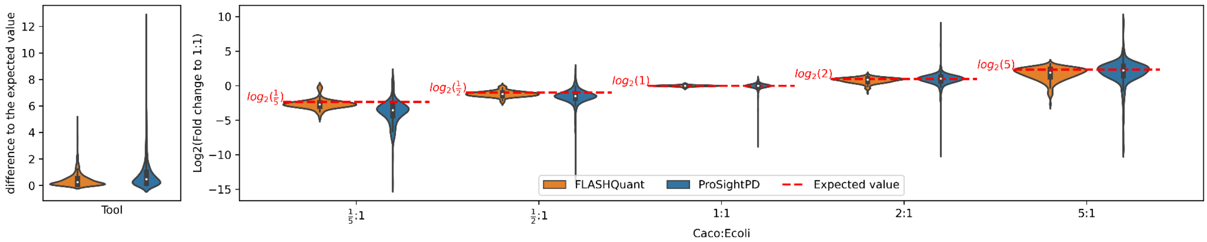


**Figure S5. Analogue of Figure 2B for the consensus feature group masses matched against Human CaCo-2 proteoform masses detected by ProSightPD Identification**

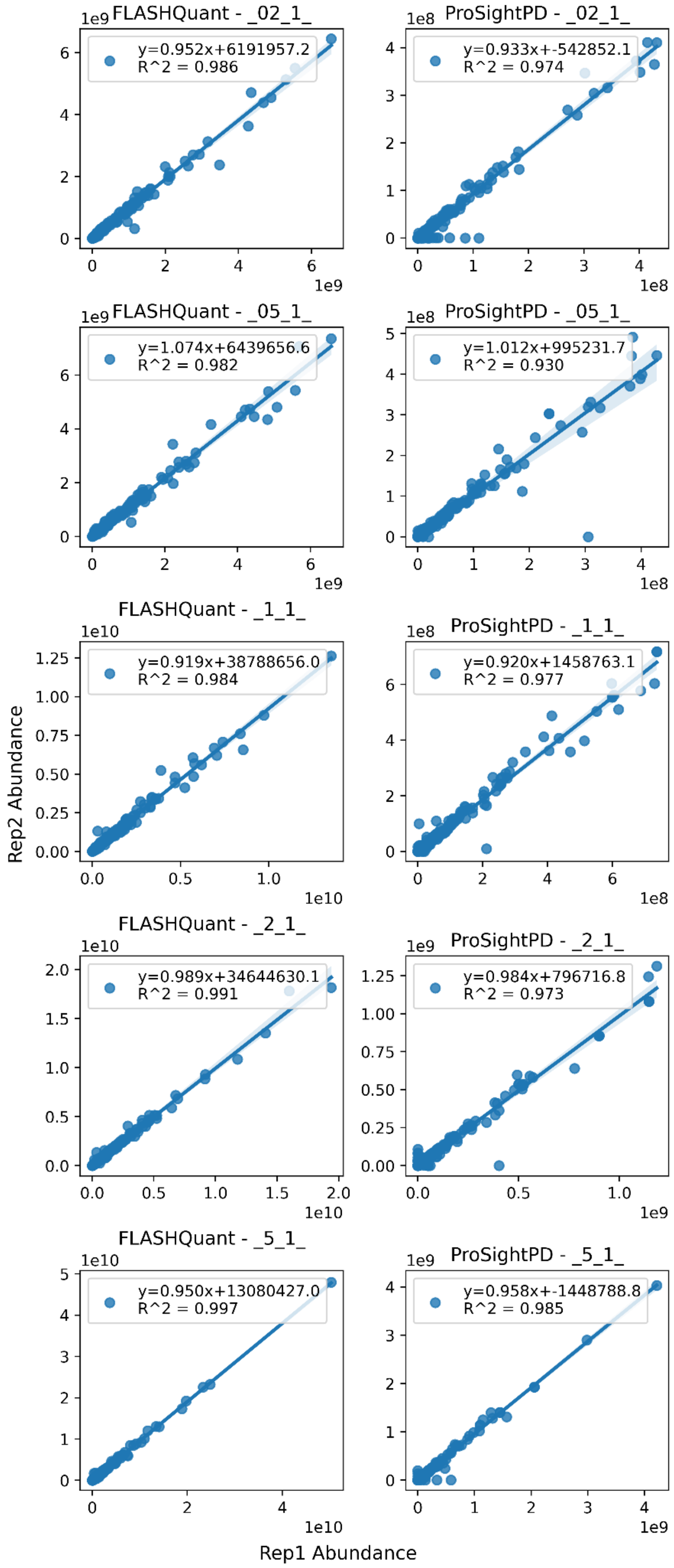


**Figure S6. Analogue of Figure S4 for the consensus feature groups matched against ProSightPD identification**

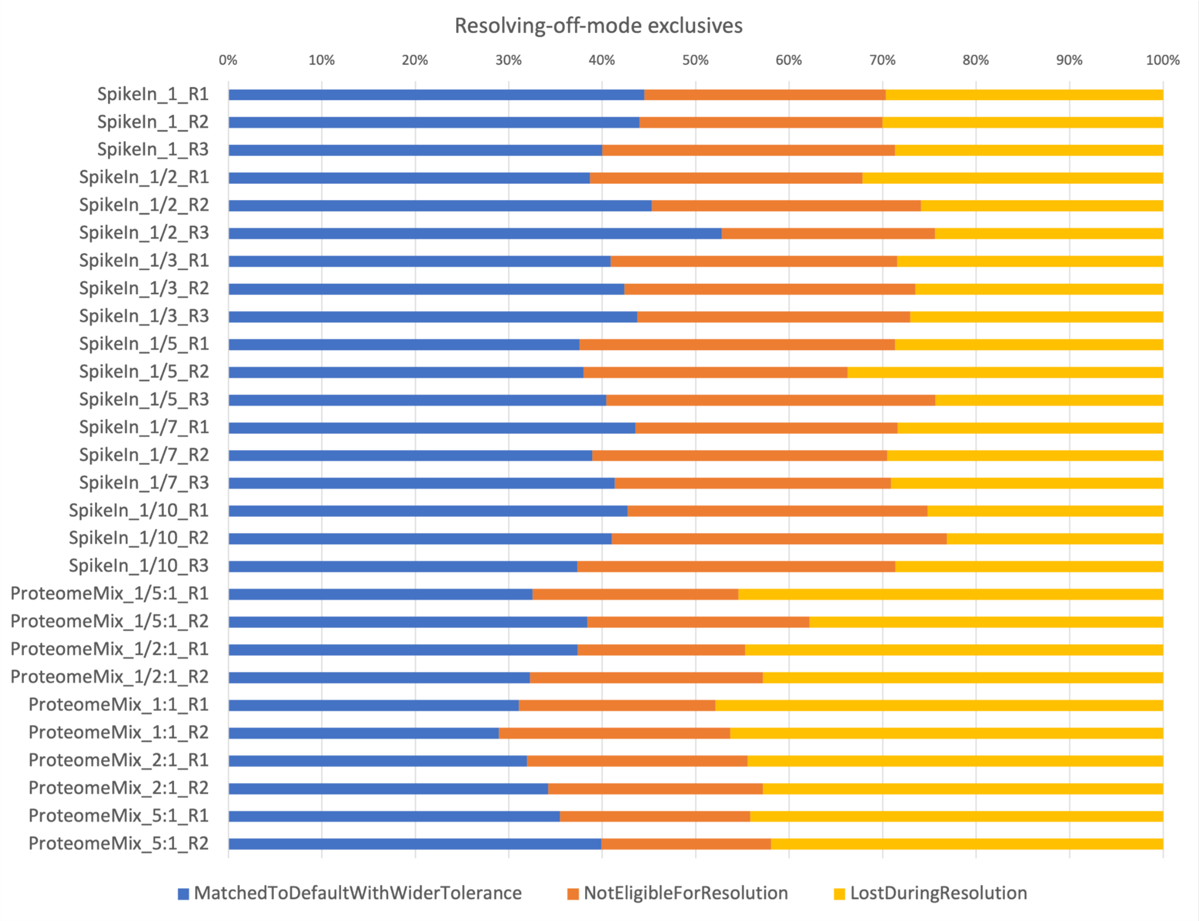


**Figure S7. Barplot of resolving-off-mode exclusives analysis**

"Resolving-off-mode" exclusives were compared with FLASHQuant results to determine where they could be originated. 30-50% of the exclusives were matched to FLASHQuant results (blue bars) when mass and retention time tolerances were widened (10 Da and 6 min). These exclusives may have been redundant and removed in FLASHQuant. Mostly, the exclusives appear to have been eliminated during the conflict resolution method (red and yellow bars). Red bars indicate that a large portion of the exclusives were not eligible for resolution (i.e., harmonics or composed of only shared m/z traces) and, thus, were removed before the resolution method. The exclusives that were likely to be removed during the conflict resolution were marked in yellow bars.

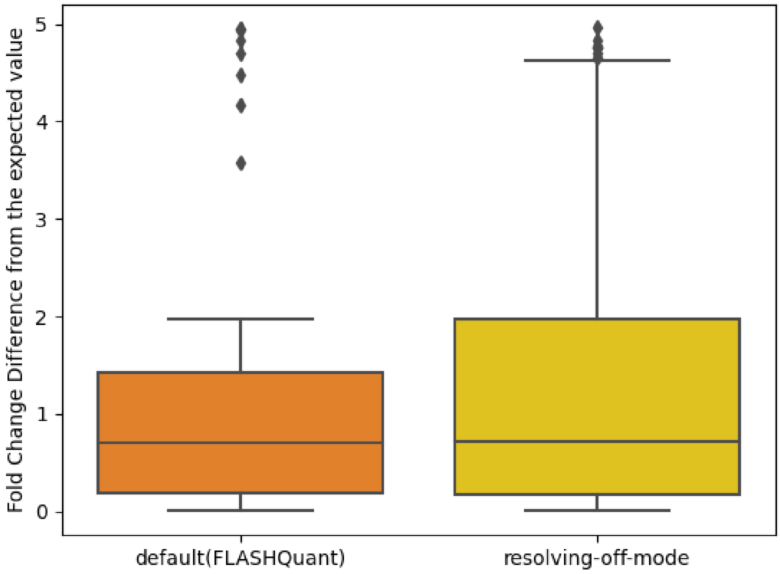


**Figure S8. Boxplot of fold change differences from the expected values comparing FLASHQuant exclusives to resolving-off-mode exclusives with the ProteomeMix dataset**

FLASHQuant exclusives show smaller fold change differences from the expected values, proving the conflict resolution method alleviated the quantification errors. With FLASHQuant and resolving-off-mode exclusives, we matched their masses to the identified proteoform masses results (of ProSightPD Search or TopPIC) to extract exclusives likely to be human Caco-2 proteoforms. Then, fold change differences of these exclusives were calculated to evaluate their quantification accuracy.

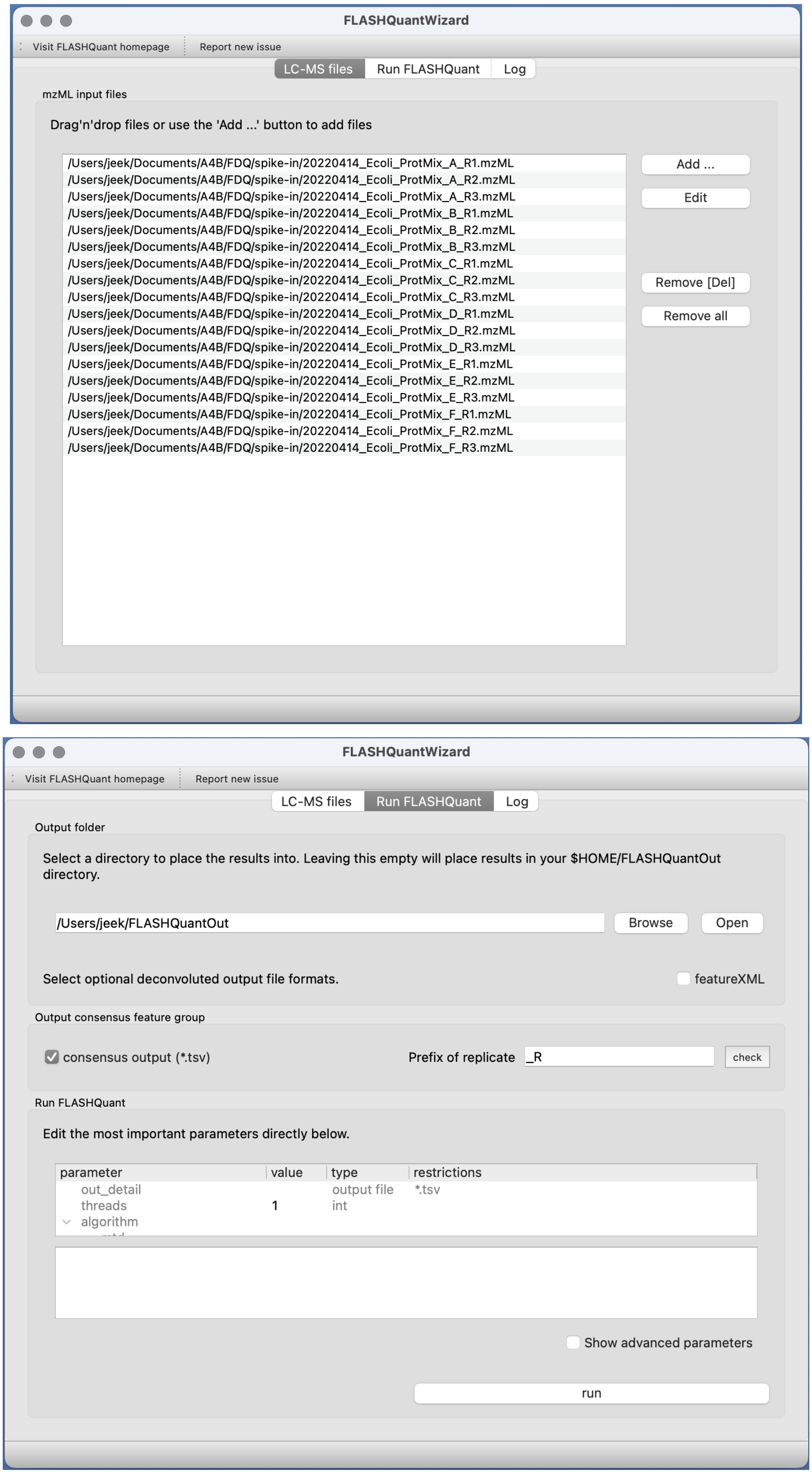


**Figure S9. FLASHQuantWizard**

Instead of a command line tool, FLASHQuant can be run with GUI. FLASHQuant will be executed per input file, and then consensus feature group detection among replicates can be optionally executed as well.

| | # Identified proteoforms |
|---|---|
| ProSightPD Search | 62 |
| TopPIC (Rep1) | 139 |
| TopPIC (Rep2) | 144 |
| TopPIC (Rep2) | 153 |

**Table S5. The number of identified proteoforms by ProSightPD Search and TopPIC for the PIPMix dataset**

This table shows the number of identified proteoforms that were used to validate consensus feature groups. The masses of all identified proteoforms (both ProSightPD Search and TopPIC) were matched to the masses of consensus feature groups within 20 ppm mass tolerance. Details about executing the identification tools are written in STAR Methods.

| Ratio (Caco-2: *E. coli*) | Caco-2 sample | *E. coli* sample | MilliQ |
|---|---|---|---|
| 5:1 | 50 µl | 10 µl | - |
| 2:1 | 20 µl | 10 µl | 30 µl |
| 1:1 | 10 µl | 10 µl | 40 µl |
| 0.5:1 | 5 µl | 10 µl | 45 µl |
| 0.2:1 | 2 µl | 10 µl | 48 µl |

**Table S6. Generation of the ProteomeMix**

The ProteomeMix data set was generated in five ratios that varied only the amount of Caco-2 protein while keeping the concentration of E. coli proteins constant.
